# SARS-CoV-2 infection suppresses ACE2 function and antiviral immune response in the upper respiratory tract of infected patients

**DOI:** 10.1101/2020.11.18.388850

**Authors:** Lucía Gutiérrez-Chamorro, Eva Riveira-Muñoz, Clara Barrios, Vanesa Palau, Marta Massanella, Edurne Garcia-Vidal, Roger Badia, Sònia Pedreño, Jordi Senserrich, Eva Rodríguez, Bonaventura Clotet, Cecilia Cabrera, Oriol Mitjà, Marta Crespo, Julio Pascual, Marta Riera, Ester Ballana

## Abstract

There is an urgent need to elucidate the molecular mechanisms underlying the transmissibility and pathogenesis of SARS-CoV-2. ACE2 is a host ectopeptidase with well-described anti-inflammatory and tissue protective functions and the receptor for the virus. Understanding SARS-CoV-2-ACE2 interaction and the expression of antiviral host genes in early infection phase is crucial for fighting the pandemic. We tested the significance of soluble ACE2 enzymatic activity longitudinally in positive nasopharyngeal swabs at two time points after symptom consultation, along with gene expression profiles of *ACE2*, its proteases, *ADAM17* and *TMPRRS2*, and interferon-stimulated genes (ISGs), *DDX58*, *CXCL10* and *IL-6*. Soluble ACE2 activity decreased during infection course, in parallel to *ACE2* gene expression. On the contrary, SARS-CoV-2 infection induced expression of the ISG genes in positive SARS-CoV-2 samples at baseline compared to negative control subjects, although this increase wanes with time. These changes positively correlated with viral load. Our results demonstrate the existence of mechanisms by which SARS-CoV-2 suppress *ACE2* expression and function casting doubt on the IFN-induced upregulation of the receptor. Moreover, we show that initial intracellular viral sensing and subsequent ISG induction is also rapidly downregulated. Overall, our results offer new insights into ACE2 dynamics and inflammatory response in the human upper respiratory tract that may contribute to understand the early antiviral host response to SARS-CoV-2 infection.

## Introduction

Angiotensin-converting enzyme 2 (ACE2) is the cellular receptor for severe acute respiratory syndrome coronavirus 2 (SARS-CoV-2), the etiological agent of the COVID-19 disease^1^. SARS-CoV-2 infection is manifested in a wide range of symptoms^2,3^, with the risk for increased disease severity correlating with male sex, advanced age and underlying comorbidities^4,5^.

One of the hallmarks of COVID-19 is a dysregulated antiviral immune response^6,7^. Interestingly, *ACE2* expression is reported to be induced upon exposure to interferon (IFN)^8^, following SARS-CoV-2 infection, in contrast with data from SARS-CoV, where cellular *ACE2* expression levels are downregulated by viral infection and linked with the pathogenicity of the virus^9^. SARS-CoV-2 may also antagonize initial viral sensing and IFN responses^10,11^, although the mechanism is still unknown and the temporal relationship between viral load and host gene expression, has not been explored *in vivo*. At gene transcription level, shotgun RNA sequencing profiles of nasopharyngeal samples from infected subjects associated IFN stimulated gene (ISGs) expression with viral load, age and sex^12^. In this context, the in-depth analysis of nasopharyngeal samples may help to decipher the intricate processes of SARS-CoV-2 infection and the host immune response. Gathering this information must be crucial to better understand how the initial phase of infection may compromise the disease progression.

ACE2 is also known to be a tissue-damage protective factor, due to its monocarboxipeptidase actions on Angiotensin (ANG)-II promoting anti-inflammatory functions through Ang-(1-7) production. These two peptides of the renin angiotensin system (RAS) are involved in the regulation of blood pressure and body fluid homeostasis^13–15^. ACE2 ectodomain is released from the cell surface by ADAM17 and TMPRSS2^16,17^carboxipeptidases, being TMPRSS2 also necessary for SARS-CoV-2 viral entry due to its function in the priming of viral S protein^16^. Interestingly, once released, the ectodomain of ACE2 keeps the catalytic activity, which is able to cleave circulating vasoactive peptides^16,17^. In the context of SARS-CoV-2 infection, it might significantly influence viral spread and pathogenesis, either by limiting new viral infections acting as a soluble decoy or alternatively, by reducing local inflammation due to its tissue protective function.

Thus, studies focused on key molecular mechanisms early after SARS-CoV-2 infection are of great value to better decipher the inflammation cascade and immune signalling in the initial phase of the disease. Moreover, the role of IFN in *ACE2* expression and shedding should be deeply evaluated because of the important implications in modulating inflammatory cytokines. Here, we evaluated the interplay between ACE2 function and host immune response in nasopharyngeal samples of a cohort of SARS-CoV-2 infected subjects, delineating how viral infection modulates both ACE2 expression and function *in vivo*, as well as, its relation with immune response and viral load.

## Results

### SARS-CoV-2 infection leads to loss of ACE2 function and expression in upper respiratory tract

Because of the central role of ACE2 receptor as the viral entry point, deciphering the ACE2 functional role is critical for the understanding of the pathophysiological changes due to SARS-CoV-2 infection. Indeed, SARS-CoV-2 viral entry might trigger cleavage of ACE2 receptor ectodomain, affecting ACE2 function and systemic release.

To gain insight into ACE2 function in viral entry and early infection events, we standardized the measurement of soluble ACE2 enzymatic activity in nasopharyngeal swabs as a surrogate marker of ACE2 function in upper respiratory tract (Supplementary fig 1). We observed significant lower ACE2 enzymatic activity levels in nasopharyngeal swabs than in serum samples, but suggesting a potential role of ACE2 activity in the upper respiratory tract in SARS-CoV-2 pathogenesis.

Thus, soluble ACE2 activity was measured in a cohort of acute infected SARS-CoV-2 subjects, confirmed by a positive PCR diagnostic test and in a group of uninfected controls, matched by age and sex. Epidemiological and clinical characteristics of the infected subjects included in the study are summarized in Table 1. SARS-CoV-2 positive subjects were longitudinally followed with a first sample at the time of the recruitment (day 0) and a second nasopharyngeal swab 3 days later. ACE2 activity did not significantly differ in SARS-CoV-2 positive compared to negative subjects, although a trend to a higher activity in SARS-CoV-2 positive samples was observed (Fig 1A). Interestingly, ACE2 activity at day 3 was significantly lower than the first sample (Fig 1A, p-value=0,0048), supporting the idea that ACE2 cleavage and release occurs upon viral entry and its function is not recovered thereafter.

**Table 1.** Baseline characteristics of index cases.

|  | All<br>N=40 | Female<br>N=29 (75.5%) | Male<br>N=11 (27.5%) |
| --- | --- | --- | --- |
| **Individual’s Characteristics** |  |  |  |
| Age (years), mean (SD) | 42.1<br>(11.3) | 40.9 (10) | 45.3 (14.1) |
| **Symptoms at baseline** |  |  |  |
| Fever, N (%) | 27 (67.5) | 18 (62.1) | 9 (81.8) |
| Cough, N (%) | 32 (80) | 24 (82.8) | 8 (72.7) |
| Dyspnea, N (%) | 6 (15) | 6 (20.7) | 0 |
| **Time from onset of symptoms to 1<sup>st</sup> sample** |  |  |  |
| Days, median (IQR) | 4 (3;6) | 5 (3;6) | 4 (4;6) |
| **LOG viral load** |  |  |  |
| RT-qPCR copies/mL, mean (SD) |  |  |  |
| Day0 | 9 (1.6) | 9 (1.7) | 9 (1.6) |
| Day3 | 6.9 (1.9) | 7.1 (2.0) | 6.4 (1.3) |

IQR: interquartile range; SD: standard deviation

**Figure 1.**
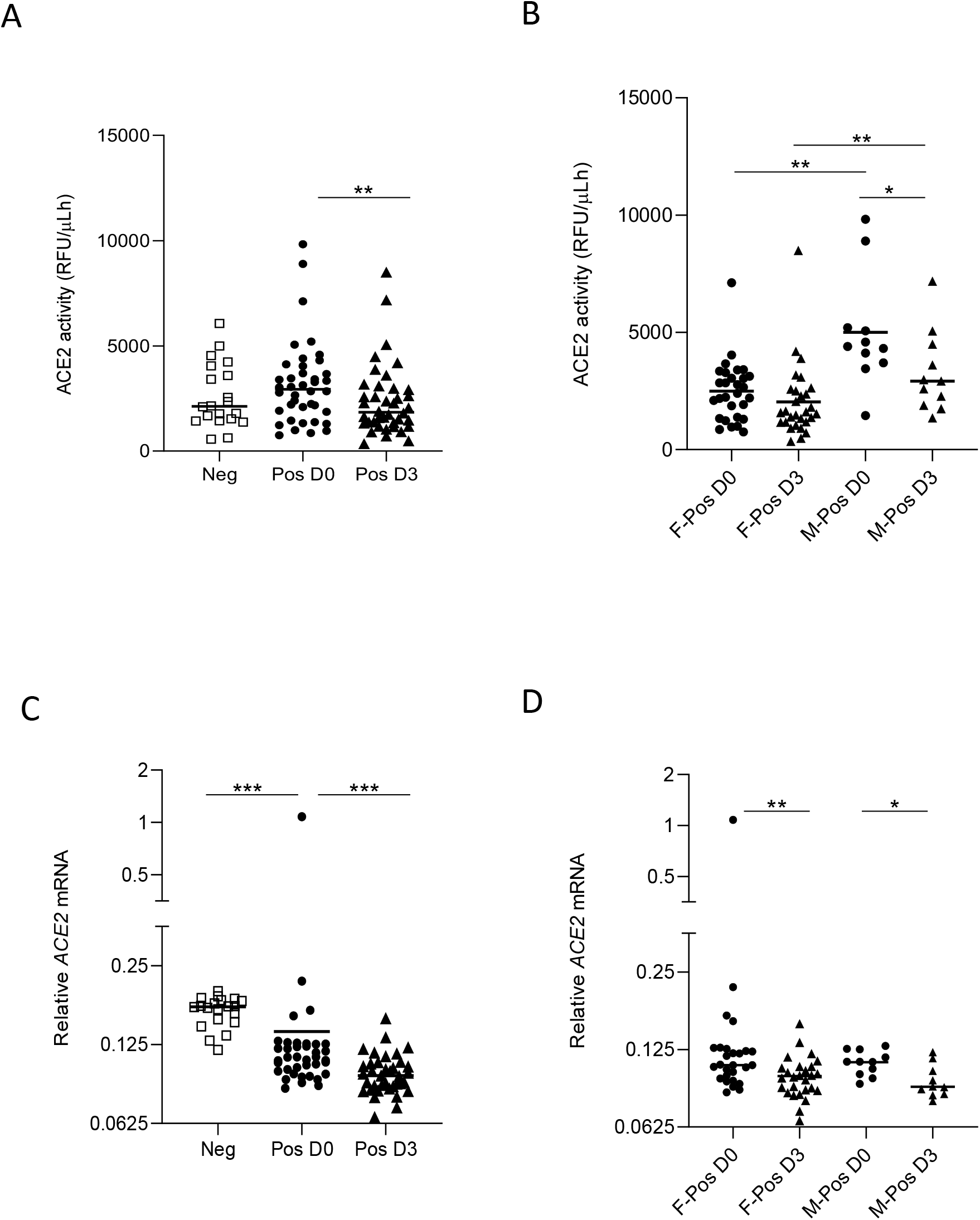
Enzymatic activity and expression of ACE2 in nasopharyngeal swabs from SARS-CoV-2 patients. **(A)** ACE2 enzymatic activity in negative (n=20, white squares, Neg) or positive SARS-CoV-2 samples (n=40), collected at the time of PCR positivity (Pos D0, black circles) and 3 days later (Pos D3, black triangles). **(B)** ACE2 activity in positive SARS-CoV-2 females (black circles, F-Pos) and males (black triangles, M-Pos) collected at the time of PCR positivity (D0) and 3 days later (D3). Data are expressed as individual relative fluorescence units (RFU) per microliter/hour. **(C)** Relative mRNA expression of *ACE2* gene in negative (n=20, white squares, Neg) or positive SARS-CoV-2 samples (n=40) at the time of PCR positivity (black circles, Pos D0) and 3 days later (black triangles, Pos D3). **(D)** Relative expression of *ACE2* in positive SARS-CoV-2 subjects stratified by sex (F-Pos, female; M-Pos, male). The mean values are presented as horizontal lines. Data was analyzed by Mann-Whitney U test, *p < 0.05, **p < 0.01, ***p < 0.001.

When samples were stratified according to sex, ACE2 activity levels were higher in SARS-CoV-2 positive males compared to females, irrespective of the day of sampling (Fig 1B, p-value= 0,0008 and p-value=0,009 for day 0 and 3, respectively), in line with previous reports showing higher plasma ACE2 levels in men^18,19^.

We next explored the temporal relationship of *ACE2* gene expression in nasopharyngeal swabs from our cohort of positive and negative SARS-CoV-2 subjects. Surprisingly, *ACE2* gene expression was significantly lower in positive SARS-CoV-2 than in negative samples (Fig 1C), in contrast to previous reports indicating IFN-dependent upregulation of *ACE2* expression upon SARS-CoV-2 infection^8^. Moreover, when comparing *ACE2* gene expression longitudinally in positive samples, *ACE2* expression showed a significant decrease overtime, following the same trend observed with ACE2 enzymatic activity, suggesting that SARS-CoV-2 is able to downregulate the expression of its receptor. No significant differences in *ACE2* gene expression were observed when samples were stratified by sex (Fig 1D).

### ACE2 function and expression correlated with SARS-CoV-2 viral load in infected subjects

SARS-CoV-2 viral load was quantified by RT-qPCR at the same timepoints were ACE2 activity and gene expression were determined, to unravel putative interactions between soluble active ACE2 and SARS-CoV-2 infection. As previously reported, viral load significantly decreased from day 0 to day 3 (mean of 2 log reduction, p-value=6,5×10^−07^, Supplementary fig 2A), showing a similar reduction to that observed for *ACE2* expression and parallel to the trend observed for ACE2 activity. No differences in viral load were identified when data was stratified by sex (Table1 and Supplementary fig 2B).

Interestingly, a significant positive correlation was found between ACE2 activity drop and viral load decrease overtime, i.e., the higher the decrease in viral load, the higher the decrease in soluble active ACE2 (spearman correlation coefficient rho=0.355, p-value=0.0246, Fig 2A). Indeed, *ACE2* gene expression also directly correlated with viral load (spearman correlation coefficient rho=0.352, p-value=0.0259 Fig 2B), strongly suggesting a direct link between viral entry and ACE2 expression and function.

**Figure 2.**
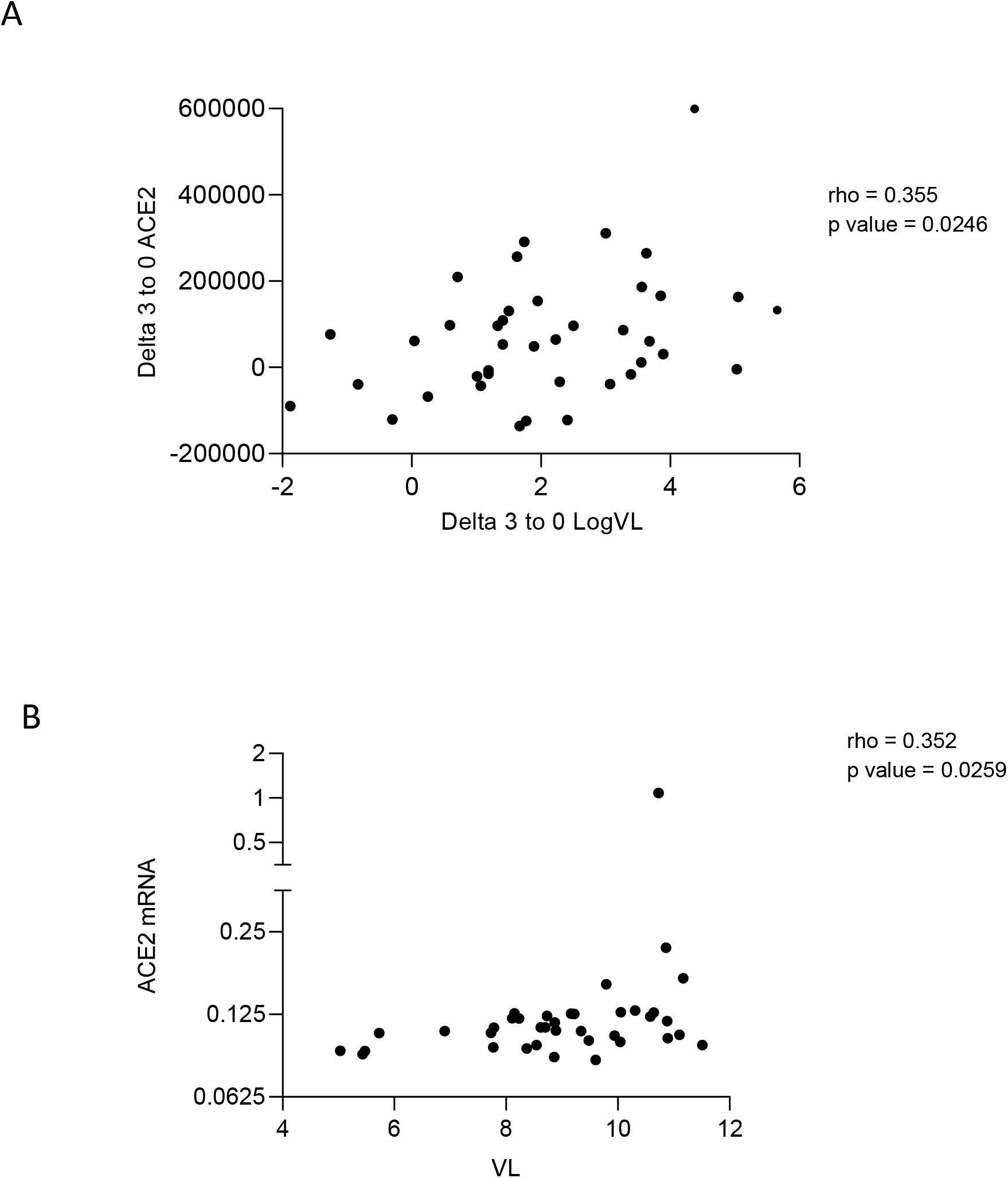
Correlation of ACE2 and SARS-CoV-2 viral load overtime. **(A)** The decrease in viral load in 3 days positively correlated with the decrease of soluble ACE2 enzymatic activity. **(B)** Viral load positively correlated with *ACE2* gene expression. Linear correlation (Spearman) r and p-values are shown.

### SARS-CoV-2 infection modulates ISG gene expression

To further explore gene expression modulation induced by SARS-CoV-2 infection, mRNA expression of *TMPRSS2* and *ADAM17*, the proteases able to cleave ACE2 ectodomain, together with distinct ISG genes were also measured. In the case of *TMPRSS2* and *ADAM17* expression, there were no significant differences in longitudinal samples obtained at different timepoints of SARS-CoV-2 positive subjects, although *TMPRSS2* expression was decreased in infected subjects compared to uninfected controls (Fig 3A, p-value<0.01).

**Figure 3.**
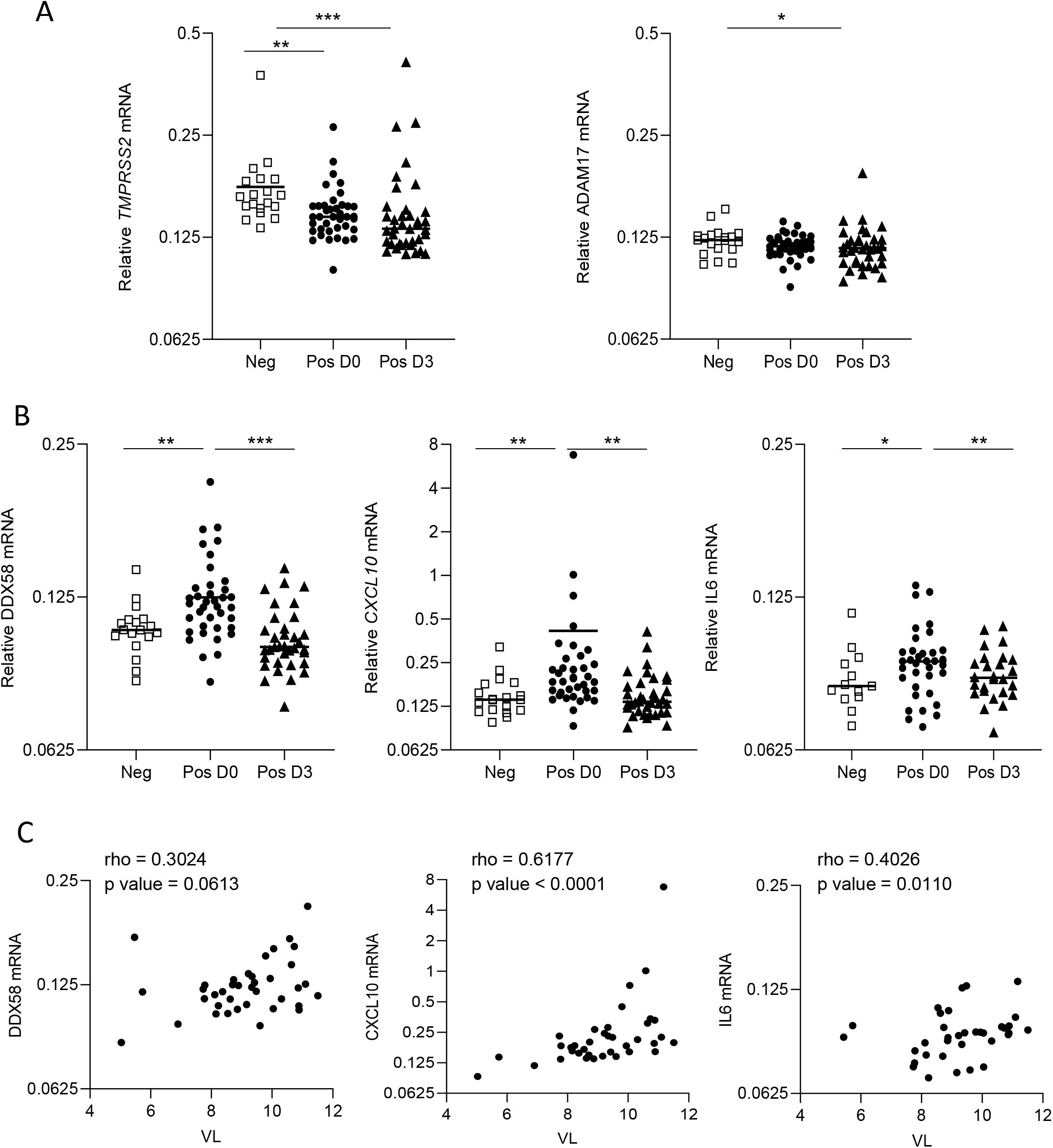
Gene expression modulation by SARS-CoV-2 in longitudinal samples of positive subjects. **(A)** Relative expression of *TMPRSS2* (left) and *ADAM17* (right) proteases in negative or positive SARS-CoV-2 longitudinal samples. **(B)** Relative expression of different ISGs. Relative mRNA expression was measured by quantitative PCR and normalized to GAPDH. Data represent 1/DCt individual values. **(C)** Positive correlations of *DDX58*, *CXCL10* and *IL6* gene expression with viral load. The mean values are presented as horizontal lines. Data was analyzed by Mann-Whitney U test, *p < 0.05, **p < 0.01, ***p < 0.001. Linear correlation (Spearman) r and p-values are shown.

On the contrary, significantly higher expression of the ISG genes, *DDX58* (RIG-I), *CXCL10* and *IL-6* genes was observed in positive SARS-CoV-2 samples at baseline compared to uninfected subjects (Fig 3B, p-value<0.05 in all cases), albeit ISG expression decreased at follow-up 3 days later, suggesting that viral infection is inducing an IFN-mediated antiviral response that is rapidly counteracted by viral proteins^10–12^.

Interestingly, increased expression of ISGs was associated with higher viral load (Fig 3C) but did not correlate with proteases expression (Supplementary Fig 3) supporting also the idea that SARS-CoV-2 infection is inducing an antiviral response upon viral entry that is afterwards suppressed to facilitate infection spread.

## Discussion

In this study, we report changes in expression of SARS-CoV-2 infection-related genes and the enzymatic activity of ACE2 in nasopharyngeal swabs contributing to understand the biology of early viral infection. Overall, our findings show the modulation induced by SARS-CoV-2 on its receptor, ACE2, at gene and soluble level throughout the first week after infection in a cohort of mild COVID-19 subjects. This modulation seems to be independent of the ISGs induced upon SARS-CoV-2 infection, also demonstrating that *ACE2* expression is not regulated by IFN. Moreover, we show that initial upregulation of antiviral immune response is directly linked to viral load and its upregulation is rapidly reversed within days during the course of infection. Our data adds key relevant knowledge to the understanding of the early antiviral host response to SARS-CoV-2 infection at the site of viral infection.

Soluble ACE2 has been described as a risk factor of death or cardiovascular events associated to several kidney or heart diseases^20–23^, and more recently, also reported in general population^19^. The role of the circulating molecule has been discussed as a feedback pathway to counterbalance the RAS and hyperinflammation activation. In our study, this scenario is also plausible in the upper airway epithelium where viral entry starts to induce the inflammatory cascade that may ultimately lead to the cytokine storm^24^. Upon infection, soluble ACE2 detection in nasopharyngeal swabs shows a slight increase in enzymatic activity, probably due to the infection itself and/or the activity of ADAM17 and/or TMPRS22 sheddases^16^. However, longitudinal evaluation of soluble ACE2 in patient-matched samples showed a significant decrease, indicating that viral infection is significantly affecting ACE2 function. Controversial observations have been published regarding ACE2 gene expression in COVID-19 infection. Previous studies on SARS-CoV linked the downregulation of ACE2 expression levels in the lungs with the pathogenicity of viral infection as a mechanism to avoid protective levels of soluble ACE2^9,25,26^. On the other hand, Ziegler and colleagues by analysis of single-cell RNA-seq datasets describe ACE2 as an IFN-stimulated gene in human epithelial tissues, upregulated following SARS-CoV-2 infection^8^. A recent publication reports the discovery of a truncated form of ACE2 gene (dACE2) as the IFN-inducible isoform of ACE2 not acting as viral receptor nor as carboxipeptidase^27^. This finding would be in line with our results showing no significant changes in ACE2 soluble activity upon SARS-CoV-2 infection, concomitant to a decrease in *ACE2* gene expression. In contrast, a clear induction of IFN stimulated genes is observed upon SARS-CoV-2 infection, also supporting the idea that full length ACE2 is not an ISG and that viral infection is indeed downregulating its expression^9^. Importantly, ACE2 expression and function correlated with viral load, further stressing the key role of ACE2 in SARS-CoV-2 pathogenesis. Indeed, age and sex differences in *ACE2* expression have also been linked to COVID-19 susceptibility and disease outcome^28^, in concordance with our data showing higher ACE2 in males compared to females.

In the setting of COVID-19 infection, soluble ACE2 has been proposed as a treatment against the spread of the virus. Clinically, changes in soluble ACE2 might imply a decrease in membrane-bound ACE2 and indicate the pathophysiological status of the patient. The presence of circulating ACE2 locally at the primary site of infection might also have implications in disease progression as a feedback pathway to counterbalance the RAS and hyperinflammation activation^29^. Parallel to the description on the biological significance of circulating ACE2, other studies showed that soluble ACE2 might be postulated as a protective molecule in some experimental models^25^ or recently as blocker for SARS-CoV-2 virus^30^. We can surmise that recombinant ACE2 might be a locally applied treatment for the first days of infection. However, further studies in extended cohorts with different infection outcomes are needed to unravel the clinical significance of soluble ACE2 in particular settings and considering each clinical situation. Although the virus-receptor interaction is needed to initiate infection, pathogenesis is a multistep process and the development of disease is influenced by multiple factors, in which target cell factors and the capacity of the host to develop a proper immune response are key. Indeed, the fact that *TMPRSS2*, but not *ADAM17* expression was downregulated upon infection, might indicate the prominent role of TMPRSS2 in the context of SARS-CoV-2 infection. On the other hand, we observed a significant upregulation of *CXCL10*, *IL-6* and *RIG-I* expression, all playing important roles in innate immune activation and clearly indicating that SARS-CoV-2 infection induces an antiviral response characterized by ISG upregulation. Importantly, the highest levels of individual ISG expression were seen in samples with the highest viral load. However, patient-matched longitudinal specimens showed a clear reduction in ISG-induced transcription 3 days later, demonstrating that the induction of an antiviral response rapidly wanes with time. In concordance with our results, it is well established the capacity of coronaviruses to manipulate immune responses and interfere with the IFN pathway, with several structural proteins (M and N) and non-structural protein (NSP1 and NSP3) from SARS-CoV and MERS-CoV acting as interferon antagonists^31^.

Collectively, our results support the existence of IFN-independent mechanisms by which SARS-CoV-2 suppress ACE2 expression and function. SARS-CoV-2 induction of ISGs at the site of infection is temporary, suggesting that the virus may also suppress intracellular viral sensing and subsequent ISG induction for favoring viral replication, although its relative contribution to disease outcome cannot be solved due to the characteristics of the studied cohort, all presenting mild forms of the disease. In depth evaluation of early changes in innate immune activation at the site of infection in patients with distinct disease severity may shed light on its putative effects on viral associated pathogenesis that may lead to different infection outcomes.

Deciphering the regulation of the ACE2 and ISG expression and function in SARS-CoV-2 target cells is a step forward in linking ACE2 levels with viral damage and COVID-19 pathology, that may help to design better strategies to efficiently clear the SARS-CoV-2 virus and minimize tissue damage. On the other hand, understanding the innate immune responses to SARS-CoV-2 and its immunoevasion approaches will improve our understanding of pathogenesis, virus clearance, and contribute toward vaccine and immunotherapeutic design and evaluation.

## Materials and methods

### Patients and samples

Nasopharyngeal swabs were obtained from 40 patients recruited under the BCN PEP CoV-2 Study (NCT04304053) during the spring SARS-CoV-2 outbreak, in Catalonia (North-East Spain). All COVID-19 cases included in the present analysis were non-hospitalized adults (i.e., ≥ 18 years of age) with quantitative PCR result available at baseline and mild symptom onset who met the diagnostic criteria for COVID-19 (fever, or acute cough, or sudden olfactory or gustatory loss, or rhinitis). Two nasopharyngeal swabs were obtained, at the time of recruitment (day 0) and at follow-up, 3 days after (day 3). Full details of the original study are reported elsewhere^32^. Similarly, a group of COVID-19 negative nasopharyngeal swabs (n=20) were collected following the same procedure and matched by age and sex to the infected patients (mean age: 39 years and 70% females). All negative samples were tested for SARS-CoV-2 infection by PCR with a negative result.

The study protocol was approved by the institutional review board of Hospital Germans Trias i Pujol. All participants provided written informed consent.

### ACE2 enzymatic assay

The ACE2 fluorescent enzymatic assay protocol was performed as previously described^23^ with slight modifications, using an ACE2-quenched fluorescent substrate (Mca-Ala-Pro-Lys(Dnp)-OH; Enzo Life Sciences). Swab samples (5μL) were incubated with ACE2 assay buffer [100mM Tris-HCl, 600mM NaCl, 10μM ZnCl2, pH 7.5 in presence of protease inhibitors 100μM captopril, 5μM amastatin, 5μM bestatin, and 10μM Z-Pro-prolinal (from Merck-Sigma-Aldrich and Enzo Life Sciences)] and 10μM fluorogenic substrate in a final volume of 100μL at 37°C for 18h. To test substrate specificity, ACE2-specific inhibitor, MLN-4760 (Merck-Sigma-Aldrich) was added. Soluble ACE2 cleaves the substrate proportionally to the enzyme activity. Measurements were carried out in duplicate for each data point. Plates (Corning clear flat-bottom 96 well black polystyrene plates) were read using a fluorescence plate reader (Ensight™ multimode plate reader, Perkin Elmer) at λ_ex_320 nm and λ_em_400 nm. Results were obtained after subtracting the background and expressed as RFU (relative fluorescent units)/μL sample/hr.

### SARS-CoV-2 PCR detection and viral load quantification

SARS-CoV-2 viral load was quantified from nasopharyngeal swabs at the distinct time points by quantitative reverse transcription PCR (qRT-PCR). RNA extraction was performed using MagMAX™ Pathogen RNA/DNA Kit (Catalog number: 4462359, ThermoFisher), optimized for a KingFisher instrument (ThermoFisher), following manufacturer’s instructions. PCR amplification was based on the 2019-Novel Coronavirus Real-Time RT-PCR Diagnostic Panel guidelines and protocol developed by the American Center for Disease control and Prevention. Briefly, a 20 μl PCR reaction was set up containing 5 μL of RNA, 1.5 μL of N3 primers and probe (2019-nCov CDC EUA Kit, #10006770, Integrated DNA Technologies) and 12.5 μl of TaqPath 1-StepRT-qPCR Master Mix (ThermoFisher). Thermal cycling was performed at 50°C for 15min for reverse transcription, followed by 95°C for 2 min and then 45 cycles of 95°C for 3 sec, 55°C for 30 sec, in the Applied Biosystems 7500 or QuantStudio5 Real-Time PCR instruments (ThermoFisher). For absolute quantification, a standard curve was built using 1/5 serial dilutions of a SARS-CoV-2 plasmid (2019-nCoV_N_Positive Control, #10006625, 200 copies/μl, Integrated DNA Technologies) and run in parallel in all PCR determinations. Viral load of each sample was determined in triplicate and mean viral load (in copies/ml) was extrapolated from the standard curve and corrected by the corresponding dilution factor.

### Quantitative RT-polymerase chain reaction (qRT-PCR)

For relative mRNA quantification RNA extraction was performed using MagMAX™ Pathogen RNA/DNA Kit, optimized for a KingFisher instrument (ThermoFisher) Reverse transcriptase was performed using the TaqPath 1-StepRT-qPCR Master Mix (ThermoFisher). mRNA relative levels of all genes were measured by two-step quantitative RT-PCR and normalized to GAPDH using the 1/DCt method. All probes were purchased from Life Technologies (TaqMan gene expression assays).

### Statistical analysis

Given that the normality was not fulfilled by the majority of variables, non-parametric methods were applied. Between group differences were evaluated trough Mann-Whitney U test. Wilcoxon signed-rank test was used to evaluate statistically significant changes in paired samples within the SARS-CoV-2 positive cohort (baseline-3 days). Correlation between variables were evaluated through Spearman’s rank correlation coefficients. Results were considered as statistically significant at p-value<0.05. Statistical analysis was performed using STATA 15.1.

## Acknowledgements

This research was supported by the crowdfunding campaign JoEmCorono (https://www.yomecorono.com/) with the contribution of over 72,000 citizens and corporations, by grants from Instituto de Salud Carlos III, Fondo de Investigación Sanitaria (FIS) PI17/00624 and PI19/00194 cofinanced by FEDER and by Grifols. EB is a research fellow from ISCIII-FIS (CPII19/00012). LGC is a research fellow from Generalitat de Catalunya AGAUR. We thank Foundation Dormeur for financial support for the acquisition of the QuantStudio-5 real time PCR system.

VP is a research fellow from ISCIII-FSE FI17/00025. CB is funded by the Catalan Health Department (Generalitat de Catalunya) contract PERIS (SLT008/18/00018). Part of this research is also supported by ISCIII-RETICS REDinREN RD16/0009/0013 and ISCIII-FIS PI16/00617. Authors also want to thank Xavier Duran for his revision and supervision of the statistical analysis.

## Author contributions

Research conceptualization and design: EB and MR. Conducted experiments: LGC, ERM, VP, RB, SP and JS. Cohort recruitment and follow up: OM and MM. Performed data analysis and interpretation: MM, CC, EGV, MR and EB. Obtained funding: CB, JP, BC, EB. Wrote or contributed to the writing and editing of the manuscript: CB, EGV, ER, CC, MC, JP, MR and EB.

