## Supplemental Figures for "SARS-CoV-2 infection suppresses ACE2 function and antiviral immune response in the upper respiratory tract of infected patients"

A

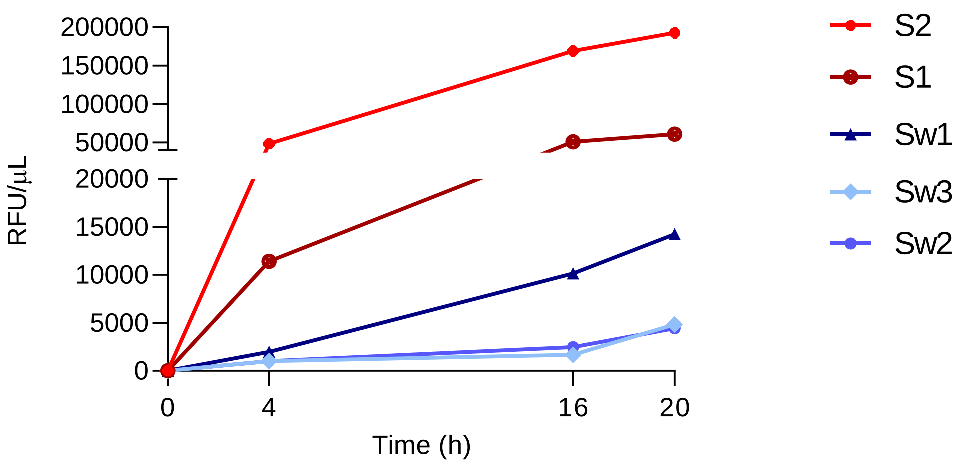


B

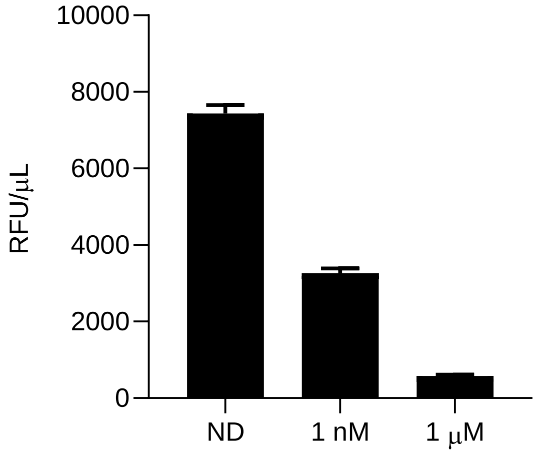


**Supplementary Figure 1. Setting up enzymatic ACE2 activity. (A)** Time course of soluble ACE2 activity in serum (S1 and S2) and nasopharyngeal swabs (Sw1, Sw2 and Sw3) samples. Results are expressed as RFU per microliter of sample. Serum samples were obtained from two healthy volunteers **(B)** ACE2 enzymatic activity in nasopharyngeal swab is blocked at concentration dependent manner by the ACE2-specific inhibitor, MLN-4760. ND; no drug. Data are shown as mean ± SEM and expressed as RFU per microliter after 18h-reaction.

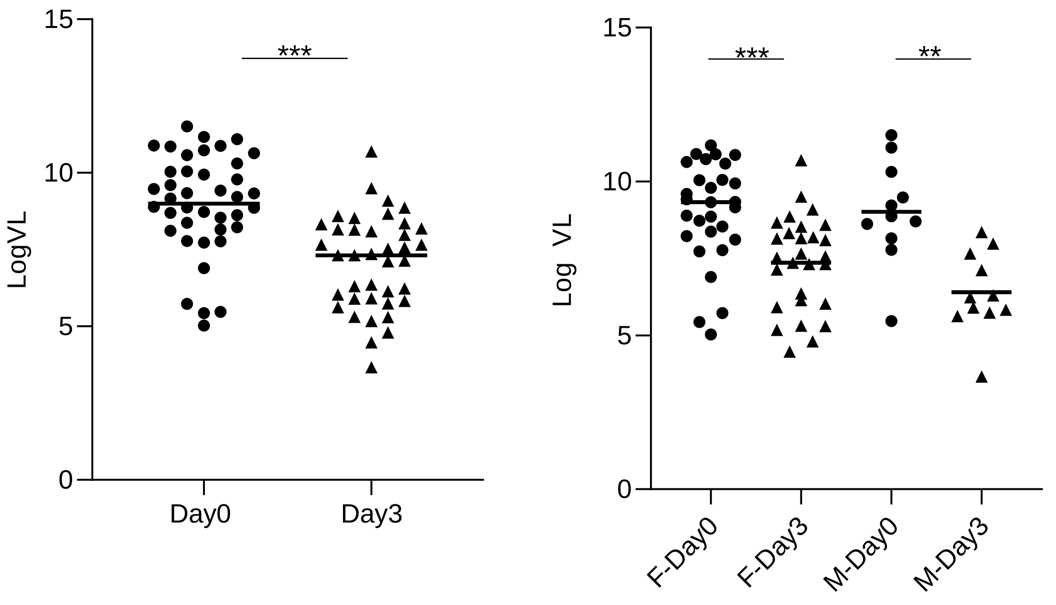


**Supplementary Figure 2. Viral load in SARS-CoV-2 positive individuals.** Left panel: Viral load at day0 and day3 of recruitment. Right panel: Sex-stratified viral load. The mean values are presented as horizontal lines. Data was analyzed by Mann-Whitney U test, *p < 0.05, **p < 0.01, ***p < 0.001.

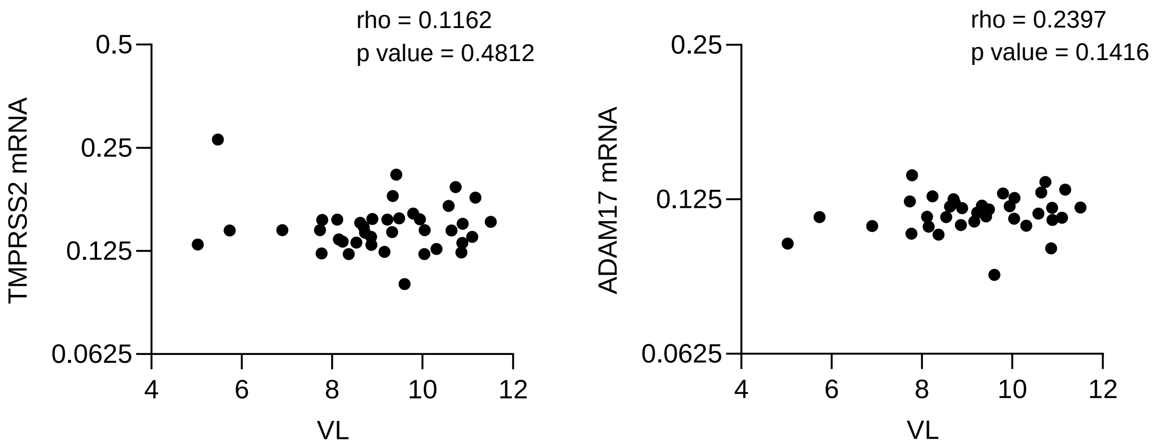


**Supplementary Figure 3. Correlation of gene expression of ACE2 shedding-related enzymes and SARS-CoV-2 viral load overtime.** Left panel: Correlation of *TMPRSS2* gene with viral load. Right panel: Correlation of *ADAM17* gene with viral load. Linear correlation (Spearman) r and p-values are shown.
